# *Candidatus* Nitrosocaldus cavascurensis, an ammonia oxidizing, extremely thermophilic archaeon with a highly mobile genome

**DOI:** 10.1101/240879

**Authors:** Sophie S. Abby, Michael Melcher, Melina Kerou, Mart Krupovic, Michaela Stieglmeier, Claudia Rossel, Kevin Pfeifer, Christa Schleper

## Abstract

Ammonia oxidizing archaea (AOA) of the phylum Thaumarchaeota are widespread in moderate environments but their occurrence and activity has also been demonstrated in hot springs. Here we present the first cultivated thermophilic representative with a sequenced genome, which allows to search for adaptive strategies and for traits that shape the evolution of Thaumarchaeota. *Candidatus* Nitrosocaldus cavascurensis has been cultivated from a hot spring in Ischia, Italy. It grows optimally at 68°C under chemolithoautotrophic conditions on ammonia or urea converting ammonia stoichiometrically into nitrite with a generation time of approximately 25h. Phylogenetic analyses based on ribosomal proteins place the organism as a sister group to all known mesophilic AOA. The 1.58 Mb genome of *Ca.* N. cavascurensis harbors an amoAXCB gene cluster encoding ammonia monooxygenase, genes for a 3-hydroxypropionate/4-hydroxybutyrate pathway for autotrophic carbon fixation, but also genes that indicate potential alternative energy metabolisms. Although a bona fide gene for nitrite reductase is missing, the organism is sensitive to NO-scavenging, underlining the importance of this compound for AOA metabolism. *Ca.* N. cavascurensis is distinct from all other AOA in its gene repertoire for replication, cell division and repair. Its genome has an impressive array of mobile genetic elements and other recently acquired gene sets, including conjugative systems, a provirus, transposons and cell appendages. Some of these elements indicate recent exchange with the environment, whereas others seem to have been domesticated and might convey crucial metabolic traits.

## Introduction

Ammonia oxidizing archaea (AOA) now collectively classified as Nitrososphaeria within the phylum Thaumarchaeota (Brochier-Armanet et al., 2008;Spang et al., 2010;Kerou, 2016) represent the sole archaeal group that is globally distributed in oxic environments efficiently competing with aerobic bacteria. Because of their large numbers in the ocean plankton, in marine sediments, in lakes and in soils, AOA are considered one of the most abundant groups of prokaryotes on this planet (Schleper et al., 2005;Prosser and Nicol, 2008;Erguder et al., 2009;Schleper and Nicol, 2010;Pester et al., 2011;Hatzenpichler, 2012;Stahl and de la Torre, 2012;Offre et al., 2013). All currently cultivated AOA strains gain energy exclusively through the oxidation of ammonia to nitrite, i.e. they perform the first step in nitrification. They grow chemolithoautotrophically from inorganic carbon supply. Some strains show growth only in the presence of small organic acids (Tourna et al., 2011;Qin et al., 2014) which seem to catalyze degradation of reactive oxygen species from the medium (Kim et al., 2016). Different from their bacterial ammonia oxidizing counterparts, AOA are often adapted to rather low levels of ammonia for growth, which seems to favor their activities in oligotrophic environments, such as the marine pelagic ocean and marine sediments, but also in acidic environments, where the concentration of ammonia decreases in favor of ammonium (Nicol et al., 2008). However, AOA occur also in large numbers in terrestrial environments, including fertilized soils and some waste water treatment plants, and several studies indicate alternative energy metabolisms (Mussmann et al., 2011;Alves et al., 2013). Although ammonia oxidation in archaea has not been biochemically resolved, the presence of genes for an ammonia monooxygenase in all AOA with remote similarity to methane and ammonia monooxygenases of bacteria implies involvement of this complex in the process (Konneke et al., 2005;Treusch et al., 2005;Nicol and Schleper, 2006). Hydroxylamine has been suggested to be the first product of ammonia oxidation (Vajrala et al., 2013), but further conversion to nitrite is performed in an unknown process, as no homolog of the bacterial hydroxylamine dehydrogenase has been found in the genomes of AOA. However, nitrous oxide (NO) has been suggested to be involved in the process, because NO production and re-consumption have been observed (Kozlowski et al., 2016b) and the NO scavenger PTIO was shown to inhibit AOA at very low concentrations (Yan et al., 2012;Shen et al., 2013;Martens-Habbena et al., 2015).

Ammonia oxidation by AOA has also been documented to occur in hot springs. With the help of molecular marker genes, diverse ammonia oxidizing thaumarchaea were found in hot terrestrial and marine environments (Reigstad et al., 2008;Zhang et al., 2008;Jiang et al., 2010;Cole et al., 2013). Furthermore, *in situ* nitrification activities up to 84°C were measured using isotopic techniques in AOA-containing habitats in Iceland (Reigstad et al., 2008), in Yellowstone National park (Dodsworth et al., 2011), in a Japanese geothermal water stream (Nishizawa et al., 2016), and in enrichment cultures from Tengchong Geothermal Field in China (Li et al., 2015). In addition, an enrichment of an ammonia oxidizing thaumarchaeon, *Ca.* Nitrosocaldus yellowstonii, from a hot spring in Yellowstone National park grew up to 74 °C confirming that archaeal ammonia oxidation, different from bacterial ammonia oxidation, indeed occurs at high temperatures (de la Torre et al., 2008).

A remarkable diversity of archaea with different metabolisms has been described from terrestrial and marine hot springs and hyperthermophilic organisms tend to be found at the base of almost all lineages of archaea. Therefore, it has often been proposed that the ancestor of all archaea and the ancestors of mesophilic lineages of archaea were hyperthermophilic or at least thermophiles (Barns et al., 1996;Lopez-Garcia et al., 2004;Gribaldo and Brochier-Armanet, 2006;Brochier-Armanet et al., 2012;Eme et al., 2013). This has been supported by sequence analyses which suggested a parallel adaptation from hot to moderate temperatures in several lineages of archaea (Groussin and Gouy, 2011;Williams et al., 2017). Indeed, the thermophilic strain *Ca.* Nitrosocaldus yellowstonii emerged in 16S rRNA phylogeny as a sister group of all known mesophilic AOA (de la Torre et al., 2008) indicating that ammonia oxidizing archaea also evolved in hot environments.

However, until now, no genome of an obligate thermophilic AOA was available to analyze in depth the phylogenetic placement of thermophilic AOA and to investigate adaptive features of these ammonia oxidizers that have a pivotal position in understanding the evolution of Thaumarchaeota.

In this work we present the physiology and first genome of an extremely thermophilic AOA of the *Nitrosocaldus* lineage that we cultivated from a hot spring in Southern Italy. We analyze its phylogeny based on ribosomal proteins and reconstruct metabolic and genomic features and adaptations.

## Material and Methods

### Sampling and enrichment

About 500 mL of mud were sampled from a terrestrial hot spring on the Italian island Ischia at “Terme di Cavascura” and stored at 4 °C until it arrived at the laboratory in Vienna (after about one week, October 2013). The temperature of 77 °C and pH 7–8 were measured *in situ* using a portable thermometer (HANNA HI935005) and pH stripes. Initial enrichments (20mL final volume) were set up in 120 mL serum flasks (two times autoclaved with MilliQ water) containing 14 mL of autoclaved freshwater medium (FWM) (de la Torre et al., 2008;Tourna et al., 2011) amended with non-chelated trace element solution (MTE) (Konneke et al., 2005), vitamin solution, FeNaEDTA (20 µL each), 2 mM bicarbonate, 1 mM ammonium chloride and 2 mL of 0.2 µm filtrated hot spring water. Serum flasks were inoculated with 4 mL (20%) of sampled mud, sealed with grey rubber stoppers (6x boiled and autoclaved) and incubated aerobically at 78 °C while rotating at 100 rpm.

### Enrichment Strategies

The temperature was changed after one week to 70 °C and medium amendment with filtrated hot spring water was stopped after four transfers as there was no effect observed. Growth was monitored by microscopy, nitrite production and ammonia consumption. Cultures were transferred at a nitrite concentration of ~700 µM and in case ammonia was depleted before the desired nitrite concentration was reached, cultures were fed with 1 mM ammonium chloride. The antibiotics Sulfamethazine, Rifampicin and Novobiocin (100 µgmL^−1^) were used alone and in combination with puruvate, glyoxylate and oxalacetate (0.1 mM), but with little effect on enrichment. Filtration of enrichments with 1.2 µm filters had no or even detrimental effect on AOA abundance when done repeatedly and 0.45 µm filtration sterilized the cultures.

As nitrite production rate increased over time, inoculum size was decreased from 20% to 10% and finally to 5%. Omitting vitamin solution from the medium led to an increase in nitrite production rate and AOA abundance. Most crucial for increasing thaumarchaeal abundance was keeping an exact timing on passaging cultures in late exponential phase (after 4 days) and setting up multiple cultures from which the best (based on microscopic observations) was used for further propagation.

### Cultivation

Cultures are routinely grown at 68 °C using 5% inoculum in 20 mL FWM amended with MTE and FeNaEDTA solutions, 1 mM NH_4_Cl, 2 mM NaHCO_3_ and are transferred every 4 days once they reach about 700 µM NO_2_^-^.

### Inhibition test with 2-phenyl-4, 4, 5, 5,-tetramethylimidazoline-1-oxyl 3-oxide (PTIO)

Cultures were grown in 20 mL FWM with 5% inoculum under standard conditions and different amounts of an aqueous PTIO stock solution were added when early to mid-exponential growth phase was reached (ca. 63 h after inoculation; 20, 100 and 400 µM final PTIO concentration). Ammonia oxidation ceased immediately at all applied PTIO concentrations, but cultures with 20 µM PTIO were able to resume nitrite production 72 h after the addition of PTIO.

### DNA Extraction

DNA was extracted from cell pellets by bead beating in pre-warmed (65 °C) extraction buffer [40.9 gL^−1^ NaCl, 12.6 gL^−1^ Na_2_SO_3_, 0.1 M Tris/HCl, 50 mM EDTA, 1% sodium dodecyl sulfate (SDS)] and phenol/ chloroform/ isoamylalcohol [25:24:1 (v/v/v), Fisher BioReagents], in a FastPrep-24 (MP) for 30 s. After centrifugation (10 min, 4 °C, 16000 g) the aqueous phase was extracted with chloroform/ isoamylalcohol [24:1 (v/v)], prior to DNA precipitation with 1 µL of glycogen (20 mg.mL^−1^) and 1 mL of PEG-solution (93.5 gL^−1^ NaCl, 300 gL^−1^ polyethylene glycol 6000) overnight at 4 °C. Nucleic acid pellets were washed, dried, resuspended in nuclease-free water and stored at −20 °C until further use (adapted from (Zhou et al., 1996;Griffiths et al., 2000)).

### Amplification of amoA gene

Specific primers were designed for the amplification of the *amo*A gene from *Candidatus* N. cavascurensis: ThAOA-277F: CCA TAY GAC TTC ATG ATA GTC and ThAOA-632R: GCA GCC CAT CKR TAN GTC CA (R. Alves and C. Schleper, unpublished). PCR conditions were 95 °C for 10 min as initialization, followed by 35 cycles of 30 s denaturing at 95 °C, 45 s primer annealing at 55 °C, and elongation at 72 °C for 45 s, finishing with 10 min final elongation at 72 °C.

### Quantitative PCR

Archaeal and bacterial 16s rRNA genes were quantified in triplicate 20 µL reactions containing 10 µL GoTaq^®^ qPCR Master Mix 2x (Promega), 0.2 mg mL^−1^ BSA and 1 µM of the appropriate primers: Arch931F [5’-AGG AAT TGG CGG GGG AGC A-3’ (Jackson et al., 2001) and Arch1100R [5’-BGG GTC TCG CTC GTT RCC-3’ (Ovreas et al., 1997)] for the archaeal 16S rRNA gene, P1 (5’-CCT ACG GGA GGC AGC AG-3’) and P2 (5’-ATT ACC GCG GCT GCT GG-3’) (Muyzer et al., 1993) for the bacterial 16S rRNA gene. Reactions were performed in a realpex^2^ (Mastercycler ep gradient S, Eppendorf AG) with the following cycling conditions: 95 °C for 2 min, followed by 40 cycles of 30 s denaturing at 95 °C, 30 s joint annealing-extension at 60 °C, and extension with fluorescence measurement at 60 °C for 30 s. Specificity of qPCR products was confirmed by melting curve analysis. Standards were prepared by cloning 16S rRNA genes from *Ca.* N. cavascurensis and *E. coli* into pGEM^®^-T Easy Vectors (Promega). These were amplified using M13 primers and concentration was determined with Qubit™ dsDNA BR Assay Kit (Thermo Fisher Scientific) before preparing serial dilutions. The efficiencies of archaeal and bacterial 16S rRNA standards were 95% and 99%, respectively.

### Fluorescence in situ hybridization

2 mL samples were centrifuged at 4°C, 16000 g for 40 min, washed in PBS-buffer and fixed with 4% paraformaldehyde for 3 h using standard protocols (Amann et al., 1990). Cells were washed two times with 1 mL PBS and finally resuspended in 200 µL of 1:1 PBS:ethanol mix before storing them at −20 °C. After dehydration in ethanol cells were hybridized overnight in hybridisation buffer with 20% formamide using probes Eub338 (Flous) 5’-GCT GCC TCC CGT AGG AGT-3’ (Amann et al., 1990) and Arch915 (dCy3) 5’-GTG CTC CCC CGC CAA TTC CT-3’ (Stahl and Amann, 1991).

### Scanning Electron Microscopy

For scanning electron microscopy, late exponential cells were harvested from 40 ml of culture by centrifugation (16000 g, 4 °C, 30 min). The cells were resuspended and washed three times (0.02 mM sodium cacodylate) prior to prefixation (0.5% glutaraldehyde in 0.02 mM sodium cacodylate) overnight, after which the concentration of glutaraldehyde was increased to 2.5% for 2h. Fixed cells were spotted onto 0.01%-poly-l-lysine coated glass slides (5 mm) and allowed to sediment for 15 min. Samples were then dehydrated using an ethanol series (30%-100%; 15 min per step) and dried by acetone (2x 100%) and HMDS (2x50%; 2x100%) substitution. The slides were subsequently placed on conductive stubs, gold coated for 30 seconds (JEOL JFC-2300HR) and analyzed (JEOL JSM-IT200).

### Genome assembly

DNA was prepared from 480 mL culture using standard procedures and sequenced using a PacBio Sequel sequencer at the VBCF (Vienna BioCenter Core Facilities GmbH). Insert size was 10 kb. Around 500000 reads were obtained with an average size of ~5 kb (N50: 6866 nt, max read length: 84504 nt).

The obtained PacBio reads were assembled using the CANU program (version 1.4, parameters “genomeSize=20m, corMhapSensitivity=normal, corOutCoverage=1000, errorRate=0.013”) (Koren et al., 2017), and then “polished” with the arrow program from the SMRT analysis software (Pacific Biosciences, USA). The *Ca.* N. cavascurensis genome consisted of two contiguous, overlapping contigs of 1580543 kb and 15533 kb. A repeated region between both extremities of the longest contig, and the 2^nd^ small contig obtained was found, and analysed using the nucmer program from the MUMMER package (see Figure Sup. 1) (Kurtz et al., 2004). This region was merged using nucmer results, and the longest contig obtained was “circularized” using the information of the nucmer program, as sequence information from the 2^nd^ contig was nearly identical (>99.5%) to that of the extremities of the long contig. The origin of replication was predicted using Ori-Finder 2 webserver (Luo et al., 2014). By analogy with *N. maritimus*, it was placed after the last annotated genomic object, before the ORB repeats and the *cdc* gene. The annotated genome sequence has been deposited into GenBank under accession number XXXXX.

### Genome dataset

Thaumarchaeota and Aigarchaeaota complete or nearly-complete genomes were downloaded from the NCBI, Genbank database, or IMG database when not available in Genbank. We included the genomes of 27 Thaumarchaeaota: Thaumarchaeota archaeon BS4 (IMG_2519899514); Thaumarchaeota group 1.1c (bin Fn1) (IMG_2558309099); Thaumarchaeota NESA-775 (Offre and Schleper, unpublished); Thaumarchaeota archaeon DS1 (IMG_2263082000); *Cenarchaeum symbiosum* A (DP000238); Thaumarchaeota archaeon CSP1 (LDXL00000000); *Nitrosopumilus adriaticus* NF5 (CP011070); Nitrosocosmicus arcticus Kfb (Alves and Schleper unpublished); *Nitrosopelagicus brevis* CN25 (CP007026); *Nitrosotenuis chungbukensis* MY2 (AVSQ00000000); *Nitrosotalea devanaterra* Nd1 (LN890280); *Nitrososphaera evergladensis* SR1 (CP007174); *Nitrosocosmicus exaquare* G61 (CP017922); *Nitrososphaera gargensis* Ga9.2 (CP002408); *Nitrosoarchaeum koreensis* MY1 (GCF_000220175); *Nitrosopumilus salaria* BD31 (GCF_000242875); *Nitrosopumilus koreensis* AR1 (CP003842); *Nitrosoarchaeum limnia* BG20 (GCF_000241145); *Nitrosoarchaeum limnia* SFB1 (CM001158); *Nitrosopumilus maritimus* SCM1 (CP000866); *Nitrosocosmicus oleophilus* MY3 (CP012850); *Nitrosopumilus piranensis* D3C (CP010868); *Nitrosopumilus sediminis* AR2 (CP003843); *Nitrosotenuis uzonensis* N4 (GCF_000723185); *Nitrososphaera viennensis* EN76 (CP007536); *Nitrosotenuis cloacae* SAT1 (CP011097); *Nitrosocaldus cavascurensis* SCU2 (this paper).

Two Aigarchaeota genomes were included: *Calditenuis aerorheumensis* (IMG_2545555825) and *Caldiarchaeum subterraneum* (BA000048). Additionally, we selected 11 Crenarchaeota genomes from Genbank to serve as an outgroup in our analysis: *Sulfolobus solfataricus* P2 (AE006641); *Pyrobaculum aerophilum* IM2 (AE009441); *Hyperthermus butylicus* DSM 5456 (CP000493); *Thermofilum pendens* Hrk 5 (CP000505); *Acidilobus saccharovorans* 345–15 (CP001742); *Desulfurococcus fermentans* DSM 16532 (CP003321); *Caldisphaera lagunensis* DSM 15908 (CP003378); *Fervidicoccus fontis* Kam940 (CP003423); *Metallosphaera sedula* DSM 5348 (CP000682); *Aeropyrum pernix* K1 (BA000002) and *Thermoproteus tenax* Kra 1 (FN869859).

### Genome annotation

The first annotation of the *Ca.* N. cavascurensis genome was obtained by the automatic annotation pipeline MicroScope (Vallenet et al., 2009;Medigue et al., 2017;Vallenet et al., 2017). Annotations from *Nitrososphaera viennensis* EN76 were carried over when aligned sequences were 35% identical (or 30% in syntenic regions) and covered 70% of the sequence. All subsequent manual annotations were held on the platform. Putative transporters were classified by screening against the Transporter Classification Database (Saier et al., 2014). To assist the annotation process, several sets of proteins of interest from Kerou et al. 2016 (EPS, metabolism), Offre et al. 2014 (transporters), Makarova et al. 2016 (archaeal pili and flagella) and ribosomal proteins (Offre et al., 2014;Kerou et al., 2016;Makarova et al., 2016) were screened specifically within our set of genomes using HMMER and HMM protein profiles obtained from databases (TIGRFAM release 15, PFAM release 28, SUPERFAMILY version 1.75), or built from arCOG families (Gough et al., 2001;Haft et al., 2003;Finn et al., 2014;Makarova et al., 2015). In the former case, the trusted cutoff “—cut_tc” was used if defined in the profiles. In the latter case, for each family of interest, sequences were extracted from the arCOG2014 database, aligned using MAFFT (“linsi” algorithm) (Katoh and Standley, 2014), and automatically trimmed at both extremities of conserved sequence blocks using a home-made script relying on blocks built by BMGE v1.12 (BLOSUM 40) (Criscuolo and Gribaldo, 2010). HMMER was then used to screen genomes for different sets of specialized families (Eddy, 2011).

In order to visualize and compare the genetic organization of sets of genes (AMO, Ure, type IV pili), we built HMM profiles (as above except alignments were manually curated for AMO and Ure), based on the corresponding arCOG families and integrated the respective sets of profiles in the MacSyFinder framework that enables the detection of sets of co-localized genes in the genome (parameters inter_gene_max_space = 10, min_nb_mandatory_genes = 1 for AMO and Ure, inter_gene_max_space = 10, min_nb_mandatory_genes = 4 for type IV pili) (Abby et al., 2014). MacSyFinder was then run with the models and profiles on the genome dataset, and the resulting JSON files were used for visualization in the MacSyView web-browser application (Abby et al., 2014). The SVG files generated by MacSyView from the predicted operons were downloaded to create figures.

### Reference phylogeny

Ribosomal proteins were identified in genomes using a set of 73 HMM profiles built for 73 arCOG families of ribosomal proteins (see Genome annotation section). When multiple hits were obtained in a genome for a ribosomal protein, the best was selected. The sequences were extracted and each family was aligned using the MAFFT program (“linsi” algorithm) (Katoh and Standley, 2014). Alignments were curated with BMGE (version 1.12, default parameters and BLOSUM 45) (Criscuolo and Gribaldo, 2010) and then checked manually. In several cases, sequences identified as part of a given ribosomal protein family did not align well and were excluded from the analysis. In the end, 59 families with more than 35 genomes represented were selected to build a phylogeny. The corresponding family alignments were concatenated, and a tree was built using the IQ-Tree program version 1.5.5 (Nguyen et al., 2015) (model test, partitioned analysis with the best model selected per gene, 1000 UF-Boot and 1000 aLRT replicates). A tree drawing was obtained with the Scriptree program (Chevenet et al., 2010), and then modified with Inkscape.

### Protein families reconstruction and phyletic patterns analysis

Homologous protein families were built for the set of genomes selected. A blast all sequences against all was performed, and the results used as input for the Silix and Hifix programs to cluster the sets of similar sequences into families (Miele et al., 2011;Miele et al., 2012). For sequences to be clustered in a same Silix family, they had to share at least 30% of identity and the blast alignment cover at least 70% of the two sequences lengths. The distribution of protein families of interest was analyzed across the genome dataset.

### Phylogenetic analyses

Sequences from Silix families of interest were extracted in fasta files, and a similarity search was performed against a large database of sequences which consisted of 5750 bacterial and archaeal genomes (NCBI Refseq, last accessed in November 2016), and the genomes from our genome dataset. Sequences with a hit having an e-value below 10^−10^ were extracted, and the dataset was then de-replicated to remove identical sequences. An alignment was then obtained for the family using MAFFT (linsi algorithm), filtered with BMGE (BLOSUM 30) and a phylogenetic tree was reconstructed by maximum-likelihood using the IQ-Tree program (version 1.5.5, best evolutionary model selected, 1000 UF-Boot and 1000 aLRT replicates). A tree drawing was obtained with the Scriptree program (Chevenet et al., 2010), and modified with Inkscape.

### Identification and annotation of integrated MGEs

Integrated mobile genetic elements were identified as described previously (Kazlauskas et al., 2017). Briefly, the integrated MGEs were identified by searching the *Ca.* N. cavascurensis genome for the presence of the hallmark genes specific to mobile genetic elements, such as integrases, casposases, VirB4 ATPases, large subunit of the terminase, major capsid proteins of achaeal viruses, etc. In the case of a positive hit, the corresponding genomic neighborhoods were analysed further. IS elements were searched for using ISfinder (Siguier et al., 2006). The precise borders of integration were defined based on the presence of direct repeats corresponding to attachment sites. The repeats were searched for using Unipro UGENE (Okonechnikov et al., 2012). Genes of integrated MGE were annotated based on PSI-BLAST searches (Altschul et al., 1997) against the non-redundant protein database at NCBI and HHpred searches against CDD, Pfam, SCOPe and PDB databases (Soding, 2005).

## Results and Discussion

### *Enrichment of Ca.* Nitrosocaldus cavascurensis

An ammonium oxidizing enrichment culture was obtained from thermal water outflow of the *Terme di Cavascura* on the Italian island Ischia. After repeated transfers into artificial freshwater medium supplemented with 1 mM ammonium and 2 mM bicarbonate, highly enriched cultures were obtained that exhibited almost stoichiometric conversion of ammonium into nitrite within 4 to 5 days (Fig. 1A). A single phylotype of AOA but no AOB or Nitrospira/commamox was identified in the enrichment via amplification and sequencing of *amo*A and 16S rRNA gene fragments (data not shown) and the presence of a single AOA phylotype was confirmed by metagenomic sequencing (see below). Its closest relative in the 16S rRNA database was fosmid 45H12 isolated from a gold mine metagenome with 99% identity and the next closest cultivated relative was *Ca.* Nitrosocaldus yellowstonii with 96% identity (Nunoura et al., 2005;de la Torre et al., 2008). The AOA has been propagated for four years within a stable enrichment culture and its relative enrichment usually ranged between 75 and 90% based on qPCR or cell counts respectively. Diverse attempts to obtain a pure culture (heat treatment, antibiotics, dilution to extinction, filtration) resulted in higher enrichments of up to 92% (based on cell counts) but not in a pure culture. In high throughput sequencing analyses using general prokaryotic primers to amplify the V2/V4 region of the 16S rRNA gene we identified phylotypes of the Deinococcus/Thermus group at up to 97% of the bacterial contaminating sequences (with 96% identity to sequences from the *Thermus* genus) and to minor extent sequences related to the Chloroflexi and Armatimonadetes (not shown). Although Aigarchaeota had been detected in earlier enrichments, the AOA was the only archaeon discovered in the high-level enrichments used in this study.

**Figure 1.**
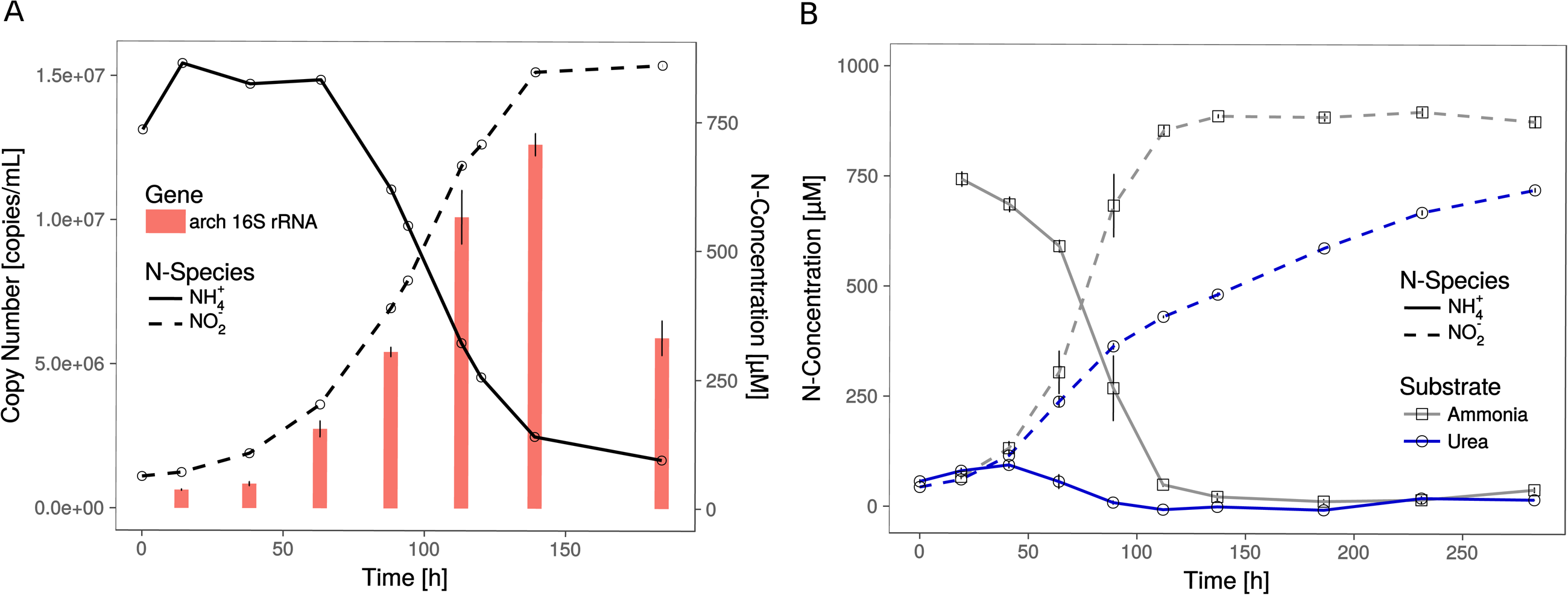
Growth of *Ca.* N. cavascurensis. **(A)** Enrichment cultures with near stoichiometric conversion of ammonia to nitrite were paralleled by cell growth, as estimated by archaeal 16S rRNA gene copies in qPCR. Gene copies declined once ammonia oxidation ceased. Data points for ammonia and nitrite are averages of biological duplicates. Technical triplicates of each biological duplicate were done to determine the gene copy number. Error bars represent the standard error of the mean. **(B)** Ammonia and successive nitrite production of enrichment cultures with 0.5 mM urea as nitrogen source. On urea as substrate ammonia accumulated during the early growth phase and the cultures showed decreasing nitrite production rate indicating growth limitation. Ammonia cultures served as control and all data points show the mean of biological triplicates with error bars representing the standard error of the mean.

The shortest generation time of the thermophilic AOA was approximately 25 h, as measured by nitrite production rates and in qPCR, which is comparable to the generation time of *Ca.* N. yellowstonii (de la Torre et al., 2008). It grew in a temperature range from about 55 to 74°C with an apparent (but relatively wide) temperature optimum around 68°C (Fig. 2).

**Figure 2.**
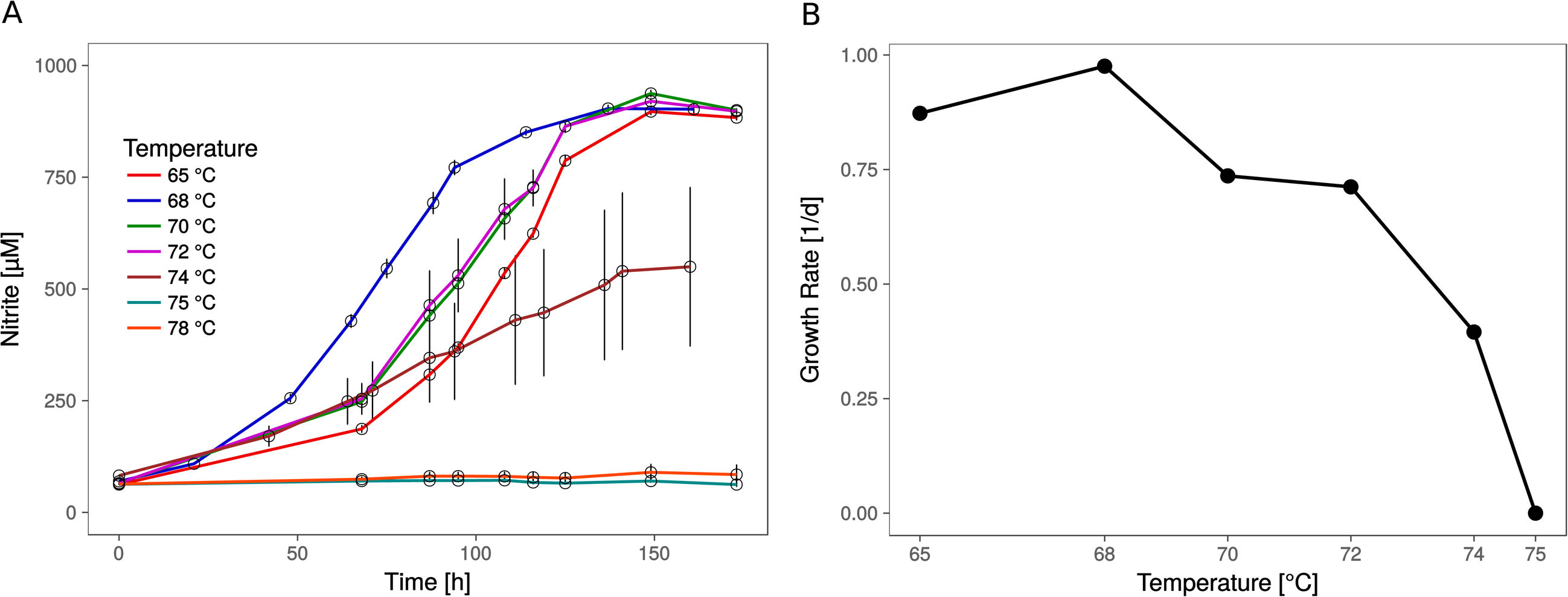
Optimal growth temperature of *Ca.* N. cavascurensis. **(A)** Nitrite production occurs within the tested temperature range (65 – 78 °C) up to 74°C, with an optimum at 68 °C. Culture quadruplets were incubated for each tested temperature and the error bars represent the standard error of the mean which increases with temperature, indicating an increase of the stochastic element within the microbial community. **(B)** Temperature dependance of growth rates showing a maximum of 0.98 d^−1^ (24.6 h generation time) at 68 °C and the highest growth temperature at 74 °C. Growth rates were calculated by linear regression of semi-logarithmically plotted nitrite values during the exponential growth phase (min. five different time points).

In fluorescence *in situ* hybridizations (FISH) with archaea-and bacteria-specific probes, all coccoid-shaped cells of less than 1 µm in diameter were assigned to the AOA, while all shorter and longer rod-shaped morphotypes were clearly assigned to bacteria (Fig. 3A, B). Scanning electron microscopy revealed the typical spherical and irregular shape of cocci, as seen e.g., for the ammonia oxidizing archaeon *Nitrososphaera viennensis* (Tourna et al., 2011) or hyperthermophilic and halophilic coccoid archaea strains with a diameter of around 0.7 µm (Fig. 3C, D). Based on its relationship with *Ca.* Nitrosocaldus yellowstonii (de la Torre et al., 2008), and the location it was sampled from (*Terme di cavascura*, Italy) we named this organism provisionally *Candidatus* Nitrosocaldus cavascurensis.

**Figure 3.**
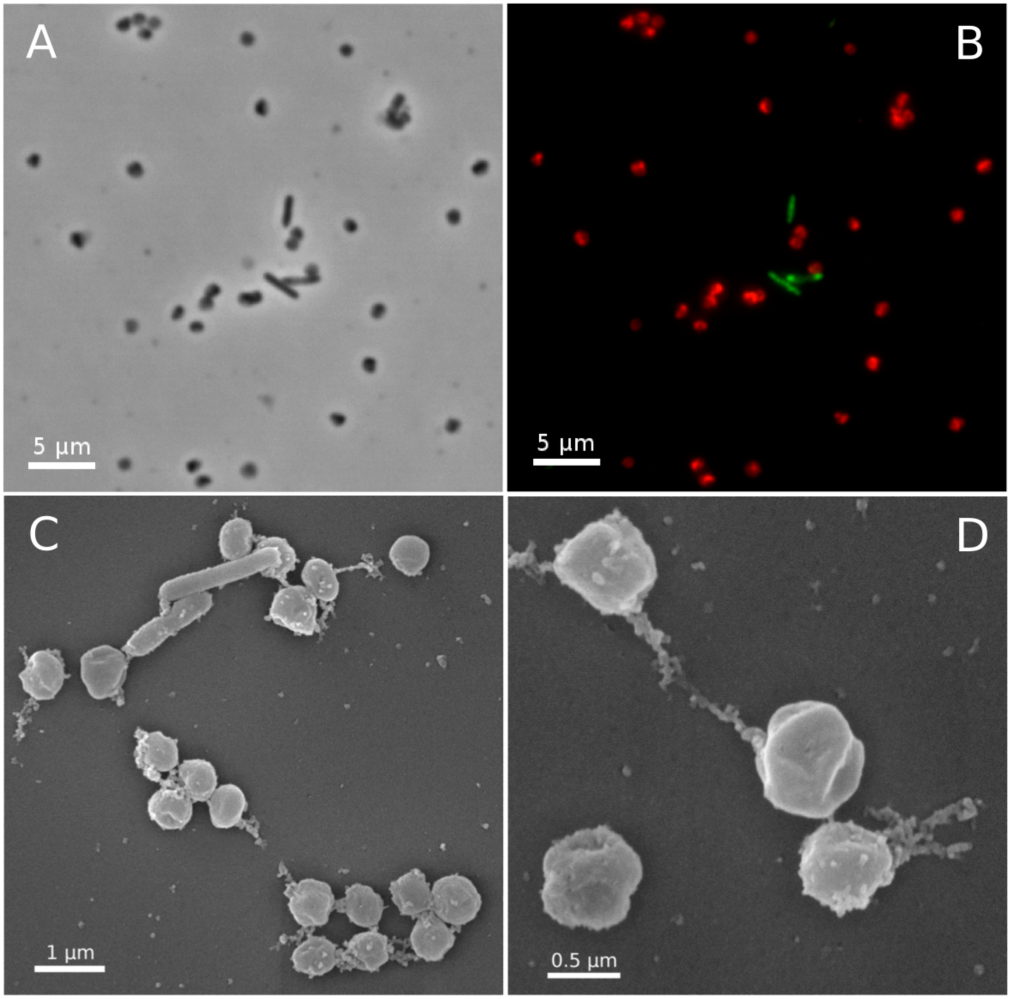
Micrographs of *Ca.* N. cavascurensis. Light **(A)** and epifluorescence micrographs **(B)** of a late exponential enrichment culture analysed with FISH. *Ca.* N. cavascurensis cells (red) and bacterial cells (green) were labeled with ARCH 915 and EUB 338 probes respectively, to give a representative picture of the enrichment state. Scanning electron micrographs **(C)** and **(D)** show the spherical nature of *Ca.* N. cavascurensis cells, having a diameter of 0.6 – 0.8 µm, small rods are visible that belong to the remaining bacterial consortium.

#### Ca. *Nitrosocaldus cavascurensis represents a deeply branching lineage of AOA*

We gathered all complete or nearly-complete 27 genome sequences available for Thaumarchaeota. Those included 23 genomes of closely-related cultivated or uncultivated AOA all harboring *amo* genes, and four genomes obtained from metagenomes that do not have genes for ammonia oxidation. Among the latter were two assembled genomes from moderate environments (Fn1 and NESA-775) and two from hot environments (BS4 and DS1) (Beam et al., 2014;Lin et al., 2015). In addition, two Aigarchaeota genomes, and a selection of representative genomes of Crenarchaeota (11) were included to serve as outgroups (Petitjean et al., 2014;Raymann et al., 2015;Adam et al., 2017;Williams et al., 2017). This resulted in a 40 genomes dataset (Material and Methods). A maximum likelihood analysis based on 59 concatenated ribosomal proteins (in one copy in at least 35 of the 40 genomes) resulted in a highly-supported tree (UF-boot and aLRT support) (Fig. 3). It confirms the monophyly of Thaumarchaeota and Nitrososphaeria (AOA), and the deeply-branching position of *Ca.* N. cavascurensis as a sister-group of all other (mesophilic) AOA, as initially indicated with a single gene (16S rRNA) phylogeny of *Ca.* Nitrosocaldus yellowstonii (de la Torre et al., 2008).

### *Genome and energy metabolism of* Ca. *Nitrosocaldus cavascurensis*

The genome of *Ca*. N. cavascurensis contains 1.58 Mbp, with 1748 predicted coding sequences, one 16S/23S rRNA operon and 29 tRNA genes. It has a G+C content of 41.6%. Similar to the genomes of most marine strains (Nitrosopumilales) and to *Ca*. Nitrosocaldus yellowstonii, but different from those of the terrestrial organisms (Nitrososphaerales) it encodes all putative subunits of the ammonia monooxygenase in a single gene cluster of the order *amoA*, *amoX*, *amoC*, *amoB* (Fig. 4) indicating that this might represent an ancestral gene order. Different from several other AOA it has a single copy of *amoC*. The genome contains a cluster of genes for the degradation of urea, including the urease subunits and two urea transporters (Fig. 4 and 5) with a similar structure to the urease locus of Nitrosophaerales. Accordingly, growth on urea, albeit slower than on ammonia, could be demonstrated (Fig. 1B). Urease loci were so far found in all Nitrososphearales genomes, and in some Nitrosopumilales (Hallam et al., 2006;Park et al., 2012;Spang et al., 2012;Bayer et al., 2016). No urease cluster could be found in non-AOA Thaumarchaeota, nor in Aigarchaeota. In Crenarchaeota, only one Sulfolobales genome harbored a urease (*Metallosphaera sedula* DSM 5348). A specific protein family of a putative chaperonin (Hsp60 homolog, arCOG01257) was found to be conserved within the urease loci of *Ca.* N. cavascurensis and the Nitrososphaerales (“soil-group” of AOA) (Fig. 4) as also observed in some bacteria (e.g., in *Haemophilus influenzae*). Unlike the conserved “thermosome” Hsp60 (also part of arCOG01257), this particular Hsp60 homolog was not found in any Nitrosopumilales genome. Given the conservation of the urease loci between *Ca.* Nitrosocaldus and Nitrosophaerales, it is possible that these genes have a common origin, and were acquired *en bloc* from bacteria by the ancestor of AOA. It is likely that the Hsp60 homolog is involved in stabilization of the urease, as demonstrated in the deltaproteobacterium *Helicobacter pylori* (Evans et al., 1992).

**Figure 4.**
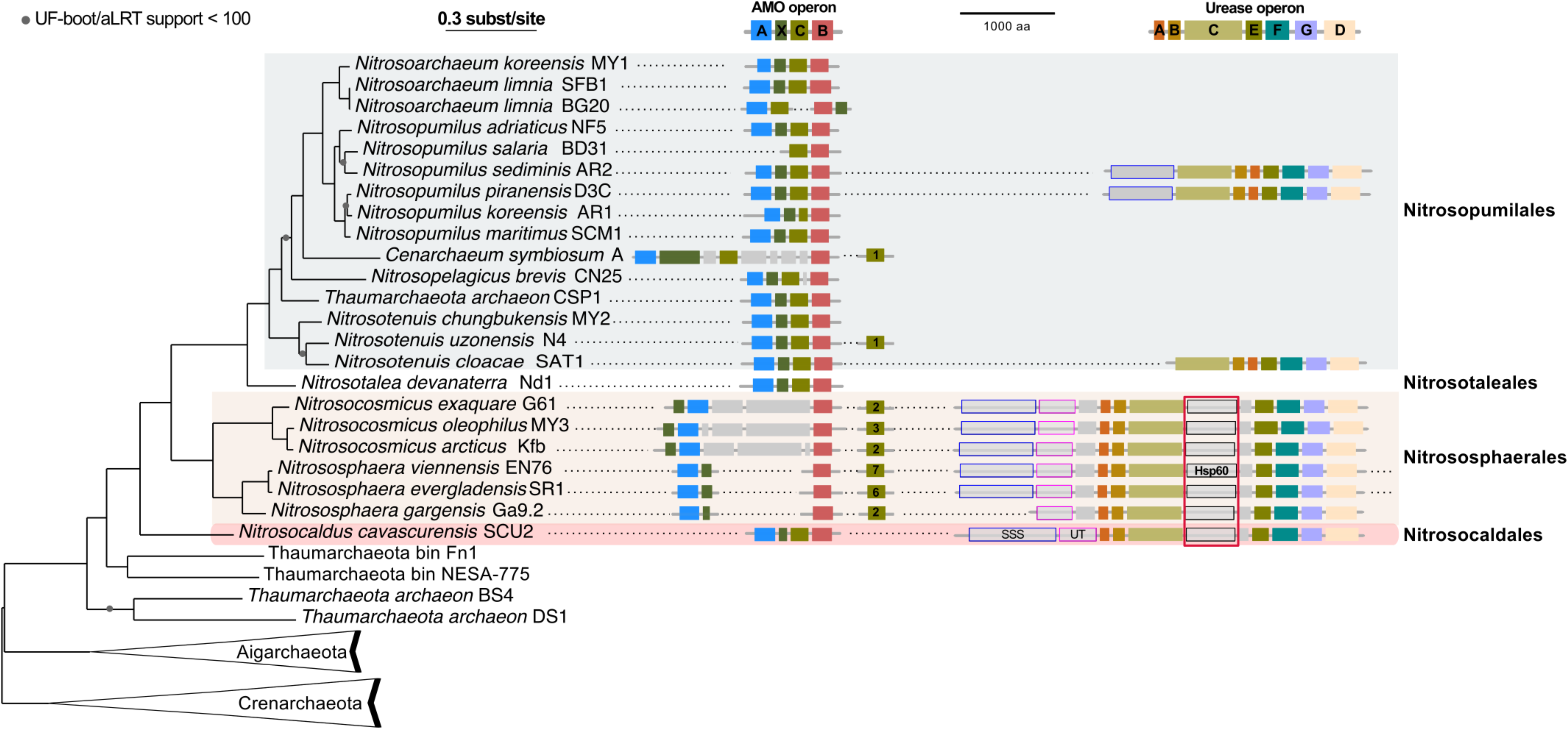
Phylogenetic position of *Ca.* N. cavascurensis, and distribution of the ammonia monooxygenase (AMO) and urease gene clusters in Thaumarchaeota genomes. This maximum-likelihood tree built with IQ-tree is based on 59 ribosomal proteins found in at least 35 genomes out of the 40 genomes dataset. AMO and Urease genes clusters found are displayed along the tree. Colors of gene boxes indicate the gene families involved in AMO or urease (*amoABCX* and *UreABCDEFG* respectively). Grey boxes correspond to other genes. Gene clusters are displayed in the orientation which maximizes the vertical alignment of the conserved genes. Genes are represented on an un-interrupted cluster when less than five genes separated them, otherwise, genes apart from more than five gene positions are displayed on another cluster. For AMO clusters, the number of different *amoC* homologs found in the genome outside of the main AMO cluster is indicated in the *amoC* box. Putative urea transporters are indicated by grey boxes outlined by blue for SSS-type urea transporter, and pink for UT urea transporter. The chaperonin Hsp60 homolog found in the urease cluster and discussed in the main text is indicated with a red box underlining its conserved position over multiple loci. Note that *N. evergladensis* and *N. viennensis* both have an extra set of three urease genes (*UreABC*) not displayed here.

**Figure 5.**
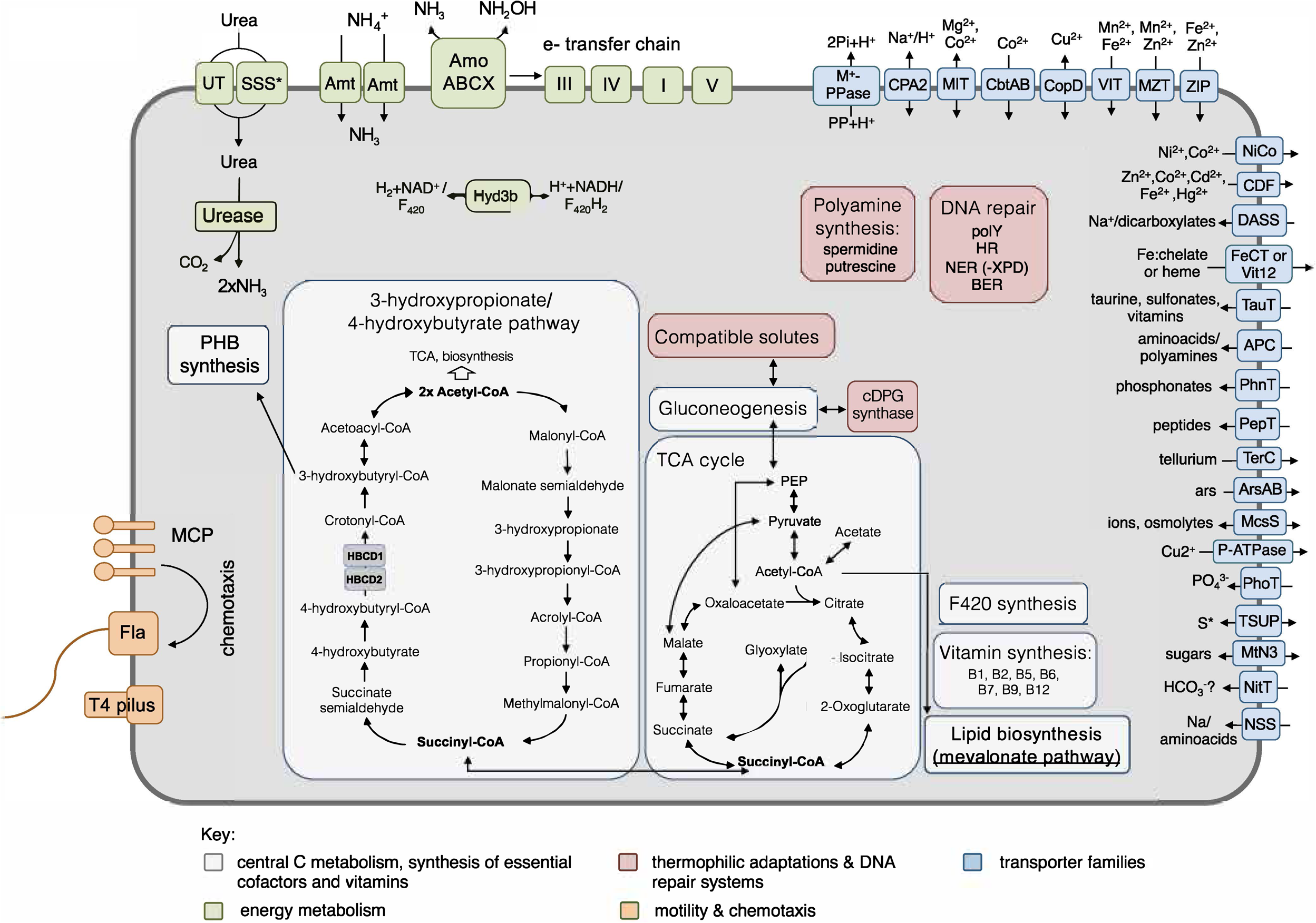
Metabolic reconstruction of *Ca*. N. cavascurensis. Schematic reconstruction of the predicted metabolic modules and other genome features of *Ca.* N. cavascurensis, as discussed in the text. Gene accession numbers and transporter classes are listed in Supplementary Table 1. HBCD: 4-hydoxybutyryl-CoA dehydratase; cDPG: cyclic 2, 3-diphosphoglycerate; MCP: methyl-accepting chemotaxis proteins; Fla: archaellum; Hyd3b: type 3b hydrogenase complex; polY: Y-family translesion polymerase; HR: homologous recombination; NER: nucleotide excision repair; BER: base excision repair; S*: organosulfur compounds, sulfite, sulfate.

Intriguingly, a gene encoding a nitrite reductase that is present in all genomes of AOA (except the sponge symbiont, *Ca.* Cenarchaeum symbiosum, (Bartossek et al., 2010)) and which is highly transcribed during ammonia oxidation in *N. viennensis* (Kerou et al., 2016) and in meta-transcriptomic datasets (Shi et al., 2011) is missing from the genome. Nitrite reductases have been postulated to be involved in ammonia oxidation by providing nitric oxide (NO) for the oxidation step to nitrite (Kozlowski et al., 2016a). We therefore tested, if the organism was affected by the NO-scavenger PTIO which inhibited at low concentrations all AOA tested so far (Martens-Habbena et al., 2015;Kozlowski et al., 2016a). Ammonia oxidation of *Ca.* N. cavascurensis was fully inhibited at concentrations as low as 20 µM, similar to those affecting *N. viennensis* and *N. maritimus* (Fig. 6). This indicates that NO is produced by an unknown nitrite reductase or perhaps by an unidentified hydroxylamine dehydrogenase (HAO), as recently shown for the HAO of the ammonia oxidizing bacterium *Nitrosomonas europaea* (Caranto and Lancaster, 2017). Alternatively, NO might be supplied through the activity of other organisms in the enrichment culture. Indeed we found a *nirK* gene in the genome of the Deinococcus species of our enrichment culture. The sensitivity to PTIO reinforces the earlier raised hypothesis that NO represents an important intermediate or cofactor in ammonia oxidation in archaea (Schleper and Nicol, 2010;Simon and Klotz, 2013;Kozlowski et al., 2016b).

**Figure 6.**
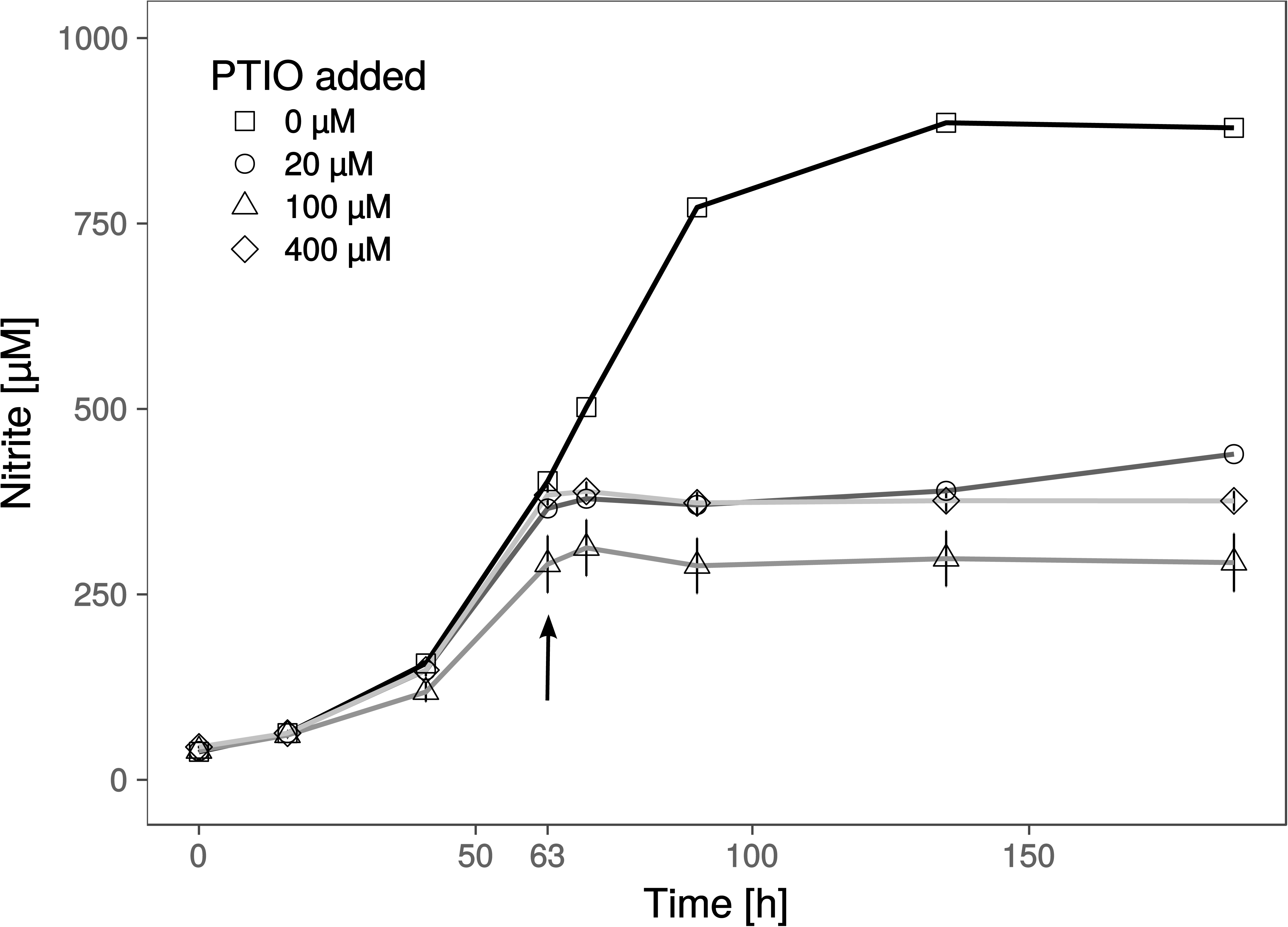
Effect of the NO scavenger 2-phenyl-4, 4, 5, 5,-tetramethylimidazoline-1-oxyl 3-oxide (PTIO) on nitrite production. Different concentrations of PTIO were added after 63 h in early to mid-exponential growth phase indicated by the arrow. Cultures with 20 µM PTIO resumed nitrite production 72 h after the addition of PTIO, while higher concentrations resulted in a complete arrest. Nitrite curves of control and 20 µM PTIO cultures show mean values from duplicates, whereas 100 and 400 µM curves are from triplicates and error bars represent the standard error of the mean.

Pathways of central carbon metabolism and -fixation of *Ca.* N. cavascurensis were found to be generally similar to those of other AOA, underlining the strikingly high similarity and conservation of metabolism within the Nitrososphaeria (Fig. 5) (Kerou et al., 2016)).

Key enzymes of the 3-hydroxypropionate/4-hydroxybutyrate carbon fixation pathway were identified in the genome of *Ca.* N. cavascurensis (supplementary table 1, (Konneke et al., 2014)). Intriguingly, and similar to *Sulfolobales*, we find two copies of the gene encoding for 4-hydoxybutyryl-CoA dehydratase located in tandem. The first copy (NCAV_0127) is homologous to the other AOA sequences which are more closely related to fermenting Clostridia rather than their crenarchaeal counterparts (“Cren Type-1”, Fig. S2), while the second copy (NCAV_0126) exhibits high similarity to genes from bacterial candidate division NC10 and Handelsmanbacteria (49% identity). In a phylogenetic tree of the protein family, it clusters together with the second group of crenarchaeal genes (“Cren Type-2”, Fig. S2, Material and Methods) which lack essential catalytic residues and which function remains unknown (Ramos-Vera et al., 2011). Könneke et al. 2014 suggested an independent emergence of the cycle in Thaumarchaeota and autotrophic Crenarchaeota based on the unrelatedness of their respective enzymes (Konneke et al., 2014). But the existence of the second gene in this deep-branching lineage rather indicates that both genes could have been present in a common ancestor of Crenarchaeota and Thaumarchaeota and that their CO_2_ fixation pathways could have a common origin. No homolog of the experimentally characterized malonic semialdehyde reductase from *N. maritimus* (Otte et al., 2015), or any other iron-containing alcohol dehydrogenase protein family member, was found in the *Ca.* N. cavascurensis genome, indicating potentially the later recruitment of this enzyme, or a case of non-orthologous gene replacement. A full oxidative TCA cycle is present in *Ca.* N. cavascurensis, including a malic enzyme and a pyruvate/phosphate dikinase connecting the cycle to gluconeogenesis. Additionally, and like most mesophilic AOA, *Ca.* N. cavascurensis encodes a class III polyhydroxyalkanoate synthase (phaEC) allowing for the production of polyhydroxyalkanoates, carbon polyester compounds that form during unbalanced growth and serve as carbon and energy reserves (Poli et al., 2011;Spang et al., 2012) (Supplementary Table 1 and Fig. 5).

The presence of a full set of genes encoding for the four subunits (and maturation factors) of a soluble type 3b [NiFe]-hydrogenase, uniquely in *Ca.* N. cavascurensis, indicates the ability to catalyze hydrogen oxidation potentially as part of the energy metabolism. This group of hydrogenases is typically found among thermophilic archaea, is oxygen-tolerant and bidirectional, and can couple oxidation of H_2_ to reduction of NAD(P) in order to provide reducing equivalents for biosynthesis, while some have been proposed to have sulfhydrogenase activity (Kanai et al., 2011;Peters et al., 2015;Greening et al., 2016) (and references therein). Although classified by sequence analysis (Sondergaard et al., 2016) and subunit composition as a type 3b hydrogenase, the alpha and delta subunits belong to the arCOG01549 and arCOG02472 families respectively, which contain coenzyme F_420_-reducing hydrogenase subunits, so far exclusively found in methanogenic archaea. Given the fact that Thaumarchaeota can synthesize this cofactor and encode a number of F_420_-dependent oxidoreductases with a yet unknown function, it is interesting to speculate whether oxidized F420 could also be a potential substrate for the hydrogenase. Expression of the hydrogenase is likely regulated through a cAMP-activated transcriptional regulator encoded within the hydrogenase gene cluster (Supplementary Table 1).

### Adaptations to thermophilic life

The molecular adaptations that enable survival and the maintenance of cell integrity at high temperatures have been the subject of intense studies since the discovery of thermophilic organisms. The issue of extensive DNA damage occurring at high temperatures has led to the study of systems of DNA stabilization and repair in thermophilic and hyperthermophilic archaea. Among them, the reverse gyrase, a type IA DNA topoisomerase shown to stabilize and protect DNA from heat-induced damage, is often (but not always (Brochier-Armanet and Forterre, 2007)) found in thermophiles, and is even considered a hallmark of hyperthermophilic organisms growing optimally above 80°C (Bouthier de la Tour et al., 1990;Forterre, 2002). However, the gene might not always be essential for survival at high temperature in the laboratory (Atomi et al., 2004;Brochier-Armanet and Forterre, 2007). Interestingly, we could not identify a gene encoding for reverse gyrase in the genome of *Ca*. N. cavascurensis. This might either reflect that its growth optimum is at the lower end of extreme thermophiles or that there is a separate evolutionary line of adaptation to thermophily possible without reverse gyrase.

The following DNA repair mechanisms were identified in the genome of *Ca.* N. cavascurensis, in agreement with the general distribution of these systems in thermophiles (for reviews see (Rouillon and White, 2011;Grasso and Tell, 2014;Ishino and Narumi, 2015)):

a) Homologous recombination repair (HR): Homologs of RadA and RadB recombinase, Mre11, Rad50, the HerA-NurA helicase/nuclease pair, and Holliday junction resolvase Hjc are encoded in the genome. These genes have been shown to be essential in other archaeal (hyper)thermophiles, leading to hypotheses regarding their putative role in replication and more generally the tight integration of repair, recombination and replication processes in (hyper)thermophilic archaea (Grogan, 2015).
b) Base excision repair (BER): the machinery responsible for the repair of deaminated bases was identified in the genome, including uracil DNA glycosylases (UDG) and putative apurinic/apyrimidinic lyases. Deletion of UDGs was shown to impair growth of (hyper)thermophilic archaea (Grogan, 2015).
c) Nucleotide excision repair (NER): Homologs of the putative DNA repair helicase: nuclease pair XPB-Bax1 and nuclease XPF were found in the genome, but repair nuclease XPD could not be identified. XPD is present in all so far analyzed mesophilic AOA, but it is also absent in other thermophiles (Kelman and White, 2005). It should be noted that NER functionality in archaea is still unclear, and deletions of the respective genes were shown to have no observable phenotype (Grogan, 2015).
d) Translesion polymerase: A Y-family polymerase with low-fidelity able to perform translesion DNA synthesis is encoded in *Ca.* N. cavascurensis. Key enzymes of all the above-mentioned systems are also found in mesophilic AOA. Given their extensive study in the crenarchaeal (hyper)thermophiles, it would be interesting to characterize their respective functions and regulation in both mesophilic and thermophilic Thaumarchaeota.
e) Bacterial-type UvrABC excision repair: In contrast to mesophilic AOA and in agreement with the known distribution of the system among mesophilic archaea and bacteria but its absence in (hyper)thermophiles, *Ca.* N. cavascurensis does not encode homologs of this repair machinery.

*Ca.* N. cavascurensis could potentially produce the polyamines putrescine and spermidine, which have been shown to bind and stabilize compacted DNA from thermal denaturation, acting synergistically with histone molecules, also present in AOA (Higashibata et al., 2000;Oshima, 2007). Although a putative spermidine synthase is also found in mesophilic AOA, the gene encoding for the previous step, S-adenosylmethionine decarboxylase (NCAV_0959), is only found in *Ca.* N. cavascurensis, located in tandem with the former. The biosynthesis of putrescine (a substrate for spermidine synthase) is unclear, since we could not identify a pyruvoyl-dependent arginine decarboxylase (ADC, catalyzing the first of the two-step biosynthesis of putrescine). However, it was shown that the crenarchaeal arginine decarboxylase evolved from an *S*-adenosylmethionine decarboxylase enzyme, raising the possibility of a promiscuous enzyme (Giles and Graham, 2008).

The production of thermoprotectant compounds with a role in stabilizing proteins from heat denaturation seems to be a preferred strategy of heat adaptation in *Ca.* N. cavascurensis. Firstly, the presence of mannosyl-3-phosphoglycerate synthase (NCAV_1295) indicates the ability to synthesize this compatible solute, shown to be involved in heat stress response and protect proteins from heat denaturation in Thermococcales (Neves et al., 2005). Homologous genes are also present in mesophilic members of the order Nitrososphaerales and other (hyper)thermophiles (Spang et al., 2012;Kerou et al., 2016). Secondly, only *Ca*. N. cavascurensis, but no other AOA encodes a cyclic 2, 3-diphosphoglycerate (cDPG) synthetase (NCAV_0908), an ATP-dependent enzyme which can synthesize cDPG from 2,3-biphosphoglycerate, an intermediate in gluconeogenesis. High intracellular concentrations of cDPG accumulate in hyperthermophilic methanogens, where it is required for the activity and thermostability of important metabolic enzymes (Shima et al., 1998).

### *Notable features of the DNA replication and cell division systems in* Ca. *N. cavascurensis*

Strikingly, only one family B replicative polymerase PolB was identified in the genome of *Ca.* N. cavascurensis (NCAV_1300), making it the only archaeon known so far to encode a single subunit of the replicative family B polymerase, as in Crenarchaeota multiple paralogs with distinct functions coexist (Makarova et al., 2014). Both subunits of the D-family polymerases PolD present in all other AOA and shown to be responsible for DNA replication in *T. kodakarensis* (also encoding both polD and polB families) (Cubonova et al., 2013) were absent from the genome, raising intriguing questions about the role of the polB family homolog in mesophilic Thaumarchaeota. Sequence analysis indicated that the *Ca.* N. cavascurensis homolog belongs to the polB1 group present exclusively in the TACK superphylum and shown recently by Yan and colleagues to be responsible for both leading and lagging strand synthesis in the crenarchaeon *Sulfolobus solfataricus* (Yan et al., 2017). However, the activity of *Sulfolobus solfataricus* PolB1 is determined based on the presence and binding of two additional proteins, PBP1 and PBP2, mitigating the strand-displacement activity during lagging strand synthesis and enhancing DNA synthesis and thermal stability of the holoenzyme, respectively. No homologs of these two additional subunits were identified in *Ca.* N. cavascurensis, raising the question of enzymatic thermal stability and efficiency of DNA synthesis on both strands.

Homologs of genes for the Cdv cell division system proteins CdvB (3 paralogues) and CdvC, but not CdvA, were identified in *Ca.* N. cavascurensis. This is surprising given that all three proteins were detected by specific immuno-labelling in the mesophilic AOA *N. maritimus*, where CdvA, CdvC and two CdvB paralogues were shown to localize mid-cell in cells with segregated nucleoids, indicating that this system mediates cell division in thaumarchaeota (Pelve et al., 2011). The *Ca.* N. cavascurensis CdvB paralogues (as all other thaumarchaeal CdvBs) all share the core ESCRT-III with the crenarchaeal CdvB sequences, contain a putative (but rather unconvincing) MIM2 motif necessary for interacting with CdvC in Crenarchaeota, while they all lack the C-terminal winged-helix domain responsible for interacting with CdvA (Samson et al., 2011;Ng et al., 2013). Interestingly, one of the *Ca.* N. cavascurensis paralogs (NCAV_0805) possesses a 40 amino-acids serine-rich C-terminal extension right after the putative MIM2 motif absent from other thaumarchaea. It is worth noting that CdvA is also absent from the published Aigarchaeota genomes *Ca.* Caldiarchaeum subterraneum (Nunoura et al., 2011) and *Ca.* Calditenuis aerorheumensis (Beam et al., 2016), while both phyla (Thaumarchaeota and Aigarchaeota) encode an atypical FtsZ homolog. Thermococcales, albeit they presumably divide with the FtsZ system, also encode CdvB and CdvC homologs, while no CdvA homolog is detectable (Makarova et al., 2010). Given the emerging differences in the molecular and regulatory aspects of the Cdv system between Crenarchaeota and Thaumarchaeota (Pelve et al., 2011;Ng et al., 2013), the intriguing additional roles of the system (Samson et al., 2017) and the fact that only CdvB and CdvC are homologous to the eukaryotic ESCRT-III system and therefore seem to have fixed roles in evolutionary terms, this observation raises interesting questions regarding the versatility of different players of the cell division apparatus.

### Ca. N. cavascurensis has a dynamic genome

The *Ca.* N. cavascurensis genome contains large clusters of genes that showed deviations from the average G+C content of the genome (*i.e.,* 41.6%), indicating that these regions might have been acquired by lateral gene transfer (Fig. 7).

**Figure 7.**
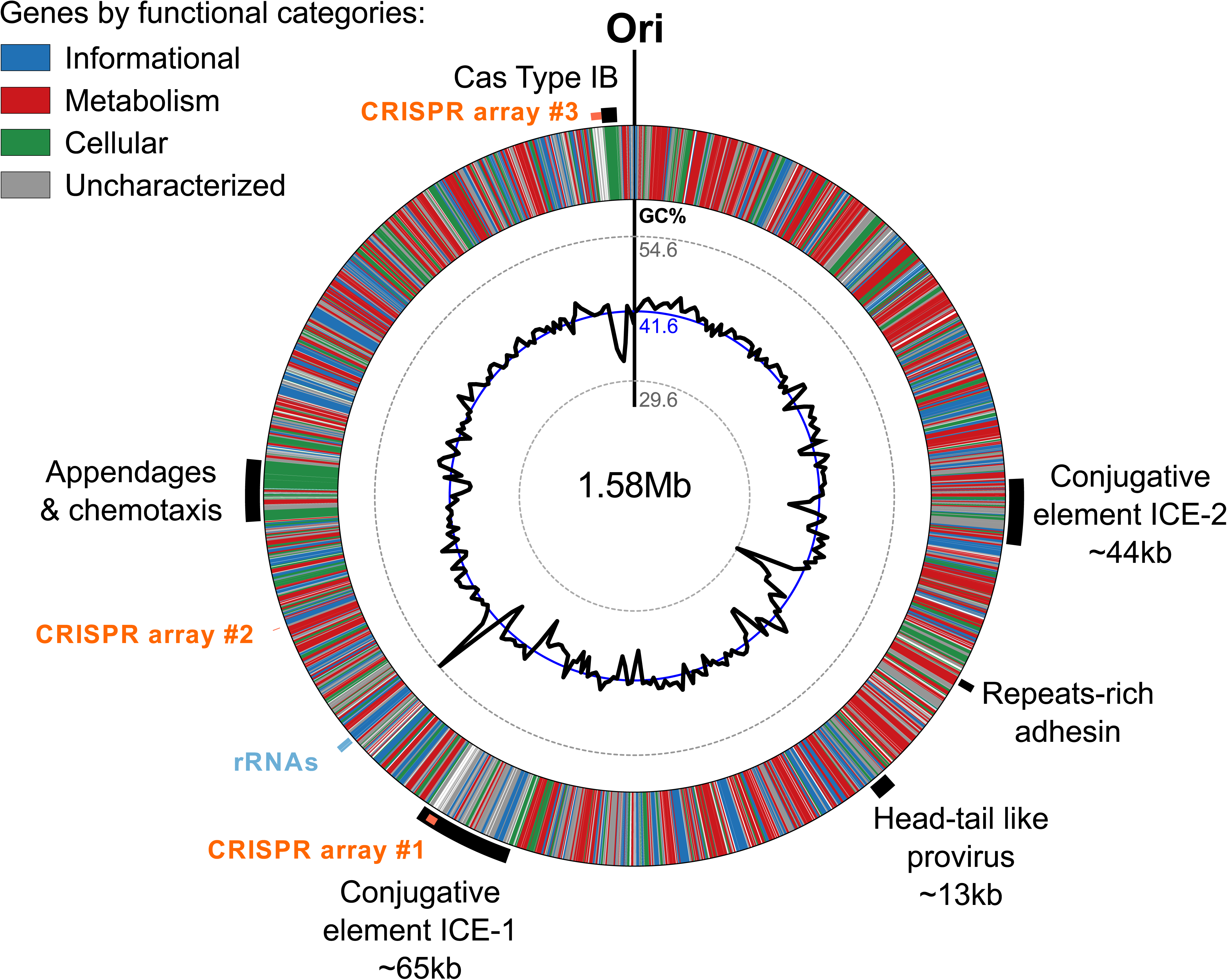
Genomic regions of atypical GC% content contain mobile genetic elements in *Ca.* N. cavascurensis. The genome is displayed in the form of a ring where genes are colored by functional categories (arCOG and COG), with the mean GC% content (blue circle) and local GC% as computed using 5kb sliding windows (black graph) in inner rings. The minimal and maximal values of local GC% are displayed as dashed grey concentric rings. The predicted origin of replication is indicated with “Ori”. Several large regions show a deviation to the average GC% that correspond to mobile genetic elements, i.e. ICEs (integrated conjugated elements), a specific pilus (see Fig. 8), and a defense system (CRISPR-Cas Type IB and three CRISPR arrays (see text).

Two of the larger regions were integrative and conjugative elements (ICE-1 and ICE-2 in Figures 7 and 8) of 65.5 kb and 43.6 kb, respectively. Both are integrated into tRNA genes and flanked by characteristic 21 bp and 24 bp-long direct repeats corresponding to the attachment sites, a typical sequence feature generated upon site-specific recombination between the cellular chromosome and a mobile genetic element (Grindley et al., 2006). ICE-1 and ICE-2 encode major proteins required for conjugation (colored in red in Fig. 8A), including VirD4-like and VirB4-like ATPases, VirB6-like transfer protein and VirB2-like pilus protein (in ICE-2). The two elements also share homologs of CopG-like proteins containing the ribbon-helix-helix DNA-binding motifs and beta-propeller-fold sialidases (the latter appears to be truncated in ICE-1). The sialidases of ICE-1 and ICE-2 are most closely related to each other (37% identity over 273 aa alignment). It is not excluded that ICE-1 and ICE-2 have inherited the conjugation machinery as well as the genes for the sialidase and the CopG-like protein from a common ancestor, but have subsequently diversified by accreting functionally diverse gene complements. Indeed, most of the genes carried by ICE-1 and ICE-2 are unrelated. ICE-1, besides encoding the conjugation proteins, carries many genes for proteins involved in DNA metabolism, including an archaeo-eukaryotic primase-superfamily 3 helicase fusion protein, Cdc6-like ATPase, various nucleases (HEPN, PD-(D/E)XK, PIN), DNA methyltransferases, diverse DNA-binding proteins and an integrase of the tyrosine recombinase superfamily (yellow in Fig. 8A). The conservation of the attachment sites and the integrase gene as well as of the conjugative genes indicates that this element is likely to be a still-active Integrative and Conjugative Element (ICE) able to integrate the chromosome and excise from it as a conjugative plasmid.

**Figure 8.**
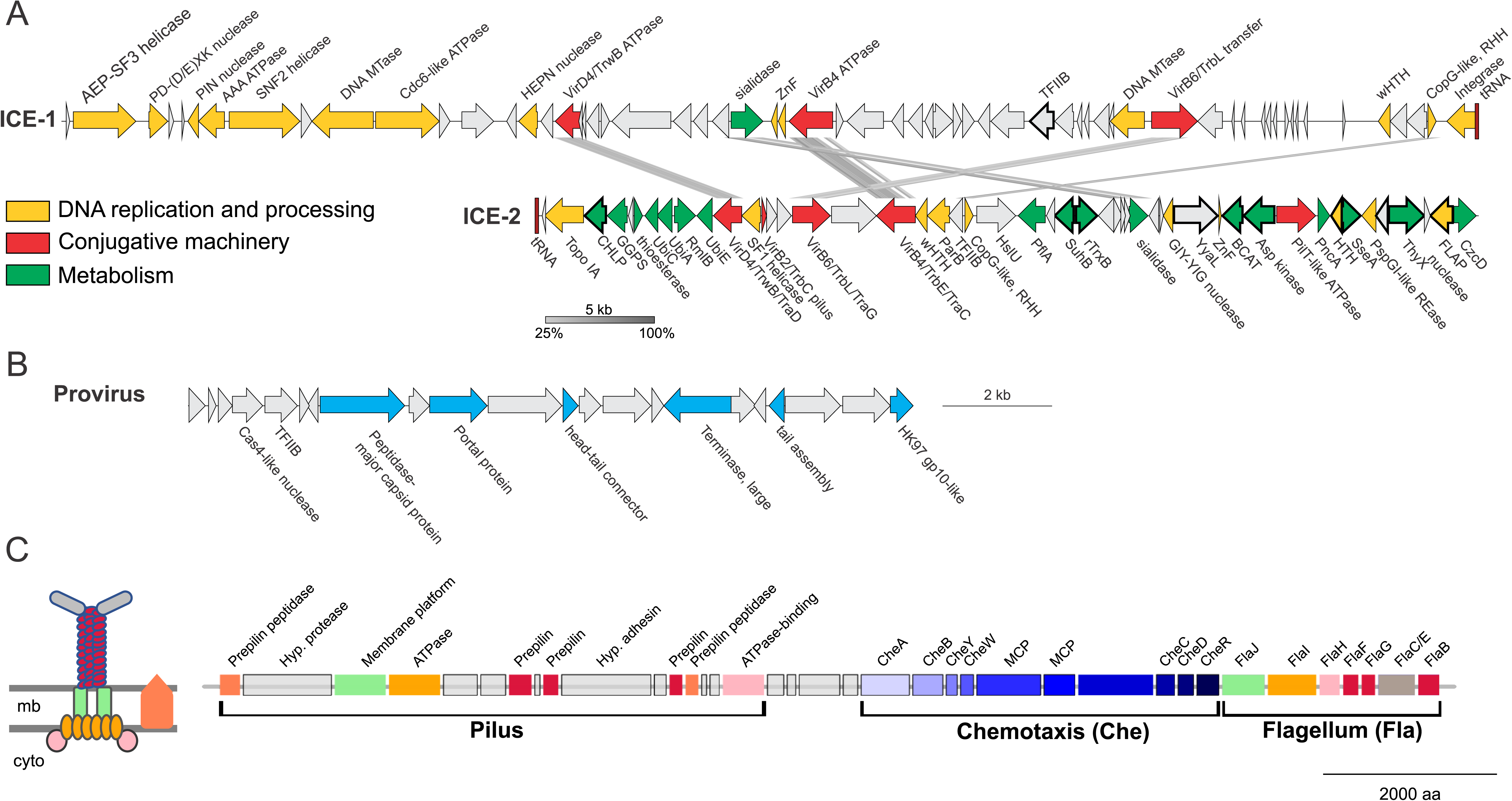
Mobile genetic elements and cell appendages encoded in the genome of *Ca.* N. cavascurensis. **(A)** Two putative Integrative Conjugative Elements (ICE) with sequence similarities shown by connexions between the two. Genes are colored by function (see legend). CHLP: geranylgeranyl diphosphate reductase; GGPS: geranylgeranyl pyrophosphate synthetase; UbiC: chorismate lyase; UbiA: 4-hydroxybenzoate octaprenyltransferase; RmlB: dTDP*glucose 4,6-dehydratase; UbiE: Ubiquinone/menaquinone biosynthesis C-methylase; HslU: ATP-dependent HslU protease; PflA: Pyruvate-formate lyase-activating enzyme; SuhB: fructose-1,6-bisphosphatase; TrxB: Thioredoxin reductase; YyaL: thoiredoxin and six-hairpin glycosidase-like domains; BCAT: branched-chain amino acid aminotransferase; PncA: Pyrazinamidase/nicotinamidase; SseA: 3-mercaptopyruvate sulfurtransferase; ThyX: Thymidylate synthase; CzcD: Co/Zn/Cd efflux system. **(B)** A head-tail like provirus from the Caudovirales order with genes in blue indicating putative viral functions. **(C)** *Ca.* N. cavascurensis specific type IV pilus locus next to a set of chemotaxis and flagellum (archaellum) genes. Homologs shared between the flagellum and the T4P are displayed with the same color. A scheme of the putative type IV pilus is shown on the left with the same color code. (inspired from drawings in (Makarova et al., 2016). mb: membrane; cyto: cytoplasm; MCP: methyl-accepting chemotaxis proteins.

ICE-2 is shorter and encodes a distinct set of DNA metabolism proteins, including topoisomerase IA, superfamily 1 helicase, ParB-like partitioning protein, GIY-YIG and FLAP nucleases as well as several DNA-binding proteins. More importantly, this element also encodes a range of proteins involved in various metabolic activities as well as a Co/Zn/Cd efflux system that might provide relevant functions to the host (green in Fig. 8A). In bacteria, ICE elements often contain cargo genes that are not related to the ICE life cycle and that confer novel phenotypes to host cells (Johnson and Grossman, 2015). It is possible that under certain conditions, genes carried by ICE-1 and ICE-2, and in particular the metabolic genes of ICE-2, improve the fitness of *Ca*. N. cavascurensis. Notably, only ICE-1 encodes an integrase, whereas ICE-2 does not, suggesting that ICE-2 is an immobilized conjugative element that can be vertically inherited in AOA. Given that the attachment sites of the two elements do not share significant sequence similarity, the possibility that ICE-2 is mobilized *in trans* by the integrase of ICE-1 appears unlikely.

The third mobile genetic element is derived from a virus, related to members of the viral order Caudovirales (Fig. 8B) (Prangishvili et al., 2017). It encodes several signature proteins of this virus group, most notably the large subunit of the terminase, the portal protein and the HK97-like major capsid protein (and several other viral homologs). All these proteins with homologs in viruses are involved in virion assembly and morphogenesis. However, no proteins involved in genome replication seem to be present. The element does not contain an integrase gene, nor is it flanked by attachment sites, which indicates that it is immobilized. Interestingly, a similar observation has been made with the potential provirus-derived element in *N. viennensis* (Krupovic et al., 2011). Given that the morphogenesis genes of the virus appear to be intact, one could speculate that these elements represent domesticated viruses akin to gene transfer agents, as observed in certain methanogenic euryarchaea (Eiserling et al., 1999;Krupovic et al., 2010), or killer particles (Bobay et al., 2014), rather than deteriorating proviruses. Notably, ICE-1, ICE-2 and the provirus-derived element all encode divergent homologs of TFIIB, a transcription factor that could alter the promoter specificity to the RNA polymerase.

A fourth set of unique, potentially transferred genes encodes a putative pilus (Fig. 8C). All genes required for the assembly of a type IV pilus (T4P) are present, including an ATPase and an ATPase binding protein, a membrane platform protein, a prepilin peptidase, and several prepilins (Makarova et al., 2016). Based on the arCOG family of the broadly conserved ATPase, ATPase-binding protein, membrane-platform protein, and prepilin peptidase (arCOG001817, -4148, -1808, and -2298), this pilus seems unique in family composition, but more similar to the archetypes of pili defined as clades 4 (A, H, I, J) by Makarova and colleagues, which are mostly found in Sulfolobales and Desulfurococcales (Fig. 4 from (Makarova et al., 2016)). Yet, the genes associated to this putative pilus seem to be more numerous, as the locus consists of approximately 16 genes (versus ~5 genes for the aap and ups in *Sulfolobus*). Such a combination of families is not found either in the T4P or flagellar loci found in analysed Thaumarchaeota genomes. Prepilins are part of the core machinery of T4P, but display a high level of sequence diversity. The three prepilins found in *Ca*. N. cavascurensis T4P locus correspond to families that are not found in any other genomes from our dataset (arCOG003872, -5987, -7276). We found a putative adhesin right in between the genes encoding the prepilins, and therefore propose that this *Ca*. N. cavascurensis-specific type IV pilus is involved in adhesion (Fig. 8C). This pilus thus appears to be unique in protein families composition when compared to experimentally validated T4P homologs in archaea (flagellum, ups, aap, bindosome (Szabo et al., 2007;Zolghadr et al., 2007;Frols et al., 2008;Tripepi et al., 2010;Henche et al., 2012)), bioinformatically predicted ones (Makarova et al., 2016), and the pili found in other Thaumarchaeota, which correspond to different types. Interestingly, this pilus gene cluster lies directly next to a conserved chemotaxis/archaellum cluster as the one found in *N. gargensis* or *N. limnia* (four predicted genes separate the T4P genes and *cheA,* Fig. 8C) (Spang et al., 2012). This suggests that this pilus might be controlled by exterior stimuli through chemotaxis. The interplay between the archaellum and pilus expression would be interesting to study in order to comprehend their respective roles.

The genome of *Ca.* N. cavascurensis also carries traces of inactivated integrase genes as well as transposons related to bacterial and archaeal IS elements, suggesting that several other types of mobile genetic elements have been colonizing the genome of *Ca*. N. cavascurensis. Collectively, these observations illuminate the flexibility of the *Ca.* N. cavascurensis genome, prone to lateral gene transfer and invasion by alien elements. Accordingly, we found a CRISPR-Cas adaptive immunity system among the sets of genes specific to *Ca*. N. cavascurensis that we could assign to the subtype I-B (Abby et al., 2014). We detected using the CRISPRFinder website (Grissa et al., 2007) at least three CRISPR arrays containing between 4 and 101 spacers presumably targeting mobile genetic elements associated with *Ca.* N. cavascurensis, reinforcing the idea of a very dynamic genome. Interestingly, the second biggest CRISPR array (96 spacers) lies within the integrated conjugative element ICE-1, which we hypothesize to be still active. This suggests that ICE-1 may serve as a vehicle for the horizontal transfer of the CRISPR spacers between *Ca*. N. cavascurensis and other organisms present in the same environment through conjugation, thus spreading the acquired immunity conferred by these spacers against common enemies.

## Conclusions

We present an obligately thermophilic ammonia oxidizing archaeon from a hot spring in the Italian island of Ischia that is related to, but also clearly distinct from *Ca.* Nitrosocaldus yellowstonii. It contains most of the genes that have been found to be conserved among AOA and are implicated in energy and central carbon metabolism, except *nirK* encoding a nitrite reductase. Its genome gives indications for alternative energy metabolism and exhibits adaptations to the extreme environment. However, it lacks an identifiable reverse gyrase, which is found in most thermophiles with optimal growth temperatures above 65°C and apparently harbors a provirus of head-and-tail structure that is usually not found at high temperature. *Ca.* N. cavascurensis differs also in its gene sets for replication and cell division, which has implications for function and evolution of these systems in archaea. In addition, its extensive mobilome and the defense system indicate that thermophilic AOA are in constant exchange with the environment and with neighboring organisms as discussed for other thermophiles (van Wolferen et al., 2013). This might have shaped and continues in shaping the evolution of thaumarchaeota in hot springs. The pivotal phylogenetic position of *Ca*. N. cavascurensis will allow reconstructing the last common ancestor of AOA and provide further insights into the evolution of this ecologically widespread group of archaea.

For our enriched strain we propose a candidate status with the following taxonomic assignment:

Nitrosocaldales order

Nitrosocaldaceae fam. and

Candidatus Nitrosocaldus cavascurensis sp. nov.

Etymology: L. adj. nitrosus, “full of natron,” here intended to mean nitrous (nitrite producer); L. masc.n. caldus, hot; cavascurensis (L.masc. gen) describes origin of sample (terme di cavascura, Ischia)

Locality: hot mud, outflow from hot underground spring, 78 °C

Diagnosis: an ammonia oxidizing archaeon growing optimally around 68 °C at neutral pH under chemolithoautotrophic conditions with ammonia or urea, spherically shaped with a diameter of approximately 0.6 to 0.8 μm, 4% sequence divergence in 16S rRNA gene from its next cultivated relative *Ca.* Nitrosocaldus yellowstonii.

## Authors contribution

CS conceived the study, MS did first enrichments, MM and CR made growth characterizations, KP did electron microscopy, SSA assembled genome and performed phylogenetic and genomic analyses for annotation, MKe annotated all metabolic and information processing genes, MKr analysed mobile genetic elements; CS, SSA, MKe wrote manuscript with contributions from MKr and MM.

## Acknowledgments

We are grateful to Lucia Monti for sharing her geological knowledge and for guiding us to the hot springs on Ischia in spring and fall 2013 and to Silvia Bulgheresi for crucial help in organising the fall expedition. We also thank Romana Bittner for excellent technical assistance and continuous cultivation and the 12 participants of the Bachelor practical class (University of Vienna) for help in sampling hot springs on Ischia and for setting up initial enrichments. We are grateful to Thomas Rattei and Florian Goldenberg for help in running computations and for providing access to the CUBE servers in Vienna, Austria. Special thanks go to LABGeM, the National Infrastructure « France Genomique » and the MicroScope team of the Genoscope (Evry, France) for providing the pipeline of annotation, and for their help during this study. We thank Eduardo Rocha and Marie Touchon for kindly providing access to their curated genome database.

## Supplementary Material

**Supplementary Figure 1. Bioinformatic genome closure in a repeat-rich region encoding a hypothetical adhesin. (A)** The two initially assembled contigs were aligned using MUMMER (Kurtz et al., 2004), revealing a sequence overlap with very high sequence identity (>99.5%) that could be resolved and resulted in the genome closure: contig 2 was a redundancy, and contig 1 was circularized through a long gene carrying repeated sequences **(B)** Analysis of this long gene showed that it encodes for an adhesin (a likely surface protein involved in adhesion). As is typical for such proteins, it contains several repeats including LamG/concavalin A domains interspersed with Ca2+-binding cadherin domains (both involved in adhesion) and terminates with two phospholipase D domains (PLD).

**Supplementary Figure 2. Phylogenetic tree of the 4-hydoxybutyryl-CoA dehydratase protein family.** A maximum likelihood phylogenetic tree was obtained with IQ-Tree v 1.5.5. The groups previously defined in (Konneke et al., 2014) “CREN type-1”, “CREN type-2”, “Anaerobe cluster”, and “AOA” are indicated by red boxes. The organisms’ class and phylum are indicated for each sequence along brackets on the right. The two homologs of this enzyme found in *Ca*. N. cavascurensis genome are indicated by a red arrow. This figure was generated using the Scriptree program (Chevenet et al., 2010), and the resulting SVG file modified using Inkscape.

**Supplementary Table 1** (supplied as an excel file). Locus tags and annotations of the main metabolic and biosynthetic pathways, information processing genes and transporters of *Ca.* Nitrosocaldus cavascurensis.

